## Supplementary material for "A conserved *C. elegans* zinc finger-homeodomain protein, ZFH-2, continuously required for structural integrity and function of alimentary tract and gonad": All Supp Figures and Tables

**for**

Sussfeld et al.

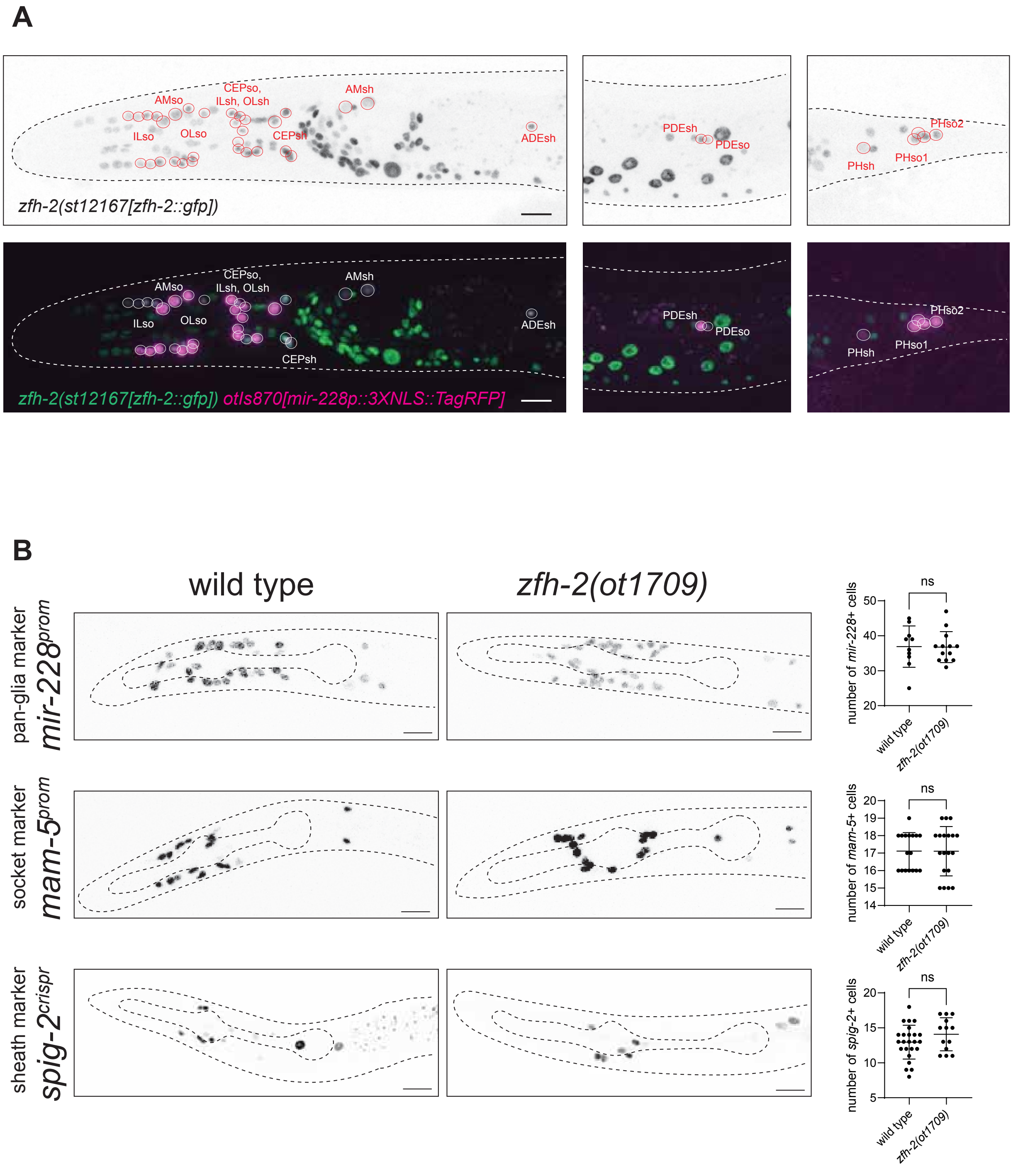


**Supplementary Figure S1**. **ZFH-2 is expressed in all ectodermal glia of *C. elegans* but is dispensable for specification of glial fates.** **(A)** A *gfp*-tagged allele of *zfh-2 (st12167[zfh-2::GFP + loxP + unc-119(+) + loxP])* reveals expression in all ectodermal glial cells, identified by the pan-glial marker *otIs870[mir-228p::3xNLS::TagRFP-T]*. **(B)** The *zfh-2* null allele *ot1709* does not significantly decrease the number of cells expressing the pan-glia marker *otIs870[mir-228p::3XNLS::TagRFP-T]* (Miska et al., 2007; Wang et al., 2024), the socket marker *otIs857[mam-5p2::3xNLS::TagRFP-T],* and the pan-sheath marker *spig-2(syb6670[spig-2::sl2::gfp::h2b])*(Aguilar and Hobert, 2024) in arrested L1 larvae. Statistical analysis was performed using unpaired t-test. ns: not significant.

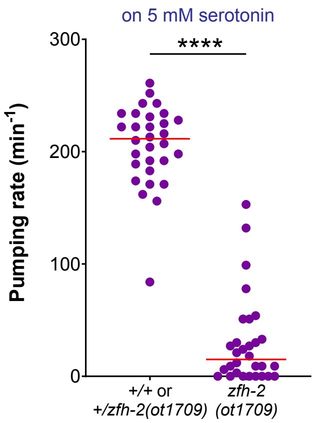


**Supplementary Figure S2: Serotonin does not rescue the pumping defects of *zfh-2* mutant.** Pharyngeal pumping rate of *zfh-2(ot1709)* L1 larval stage animals in the presence of 5 mM serotonin. Horizontal line in the middle of data points represents median value of biological replicates. **** $P < 0.0001$ in Mann-Whitney test.

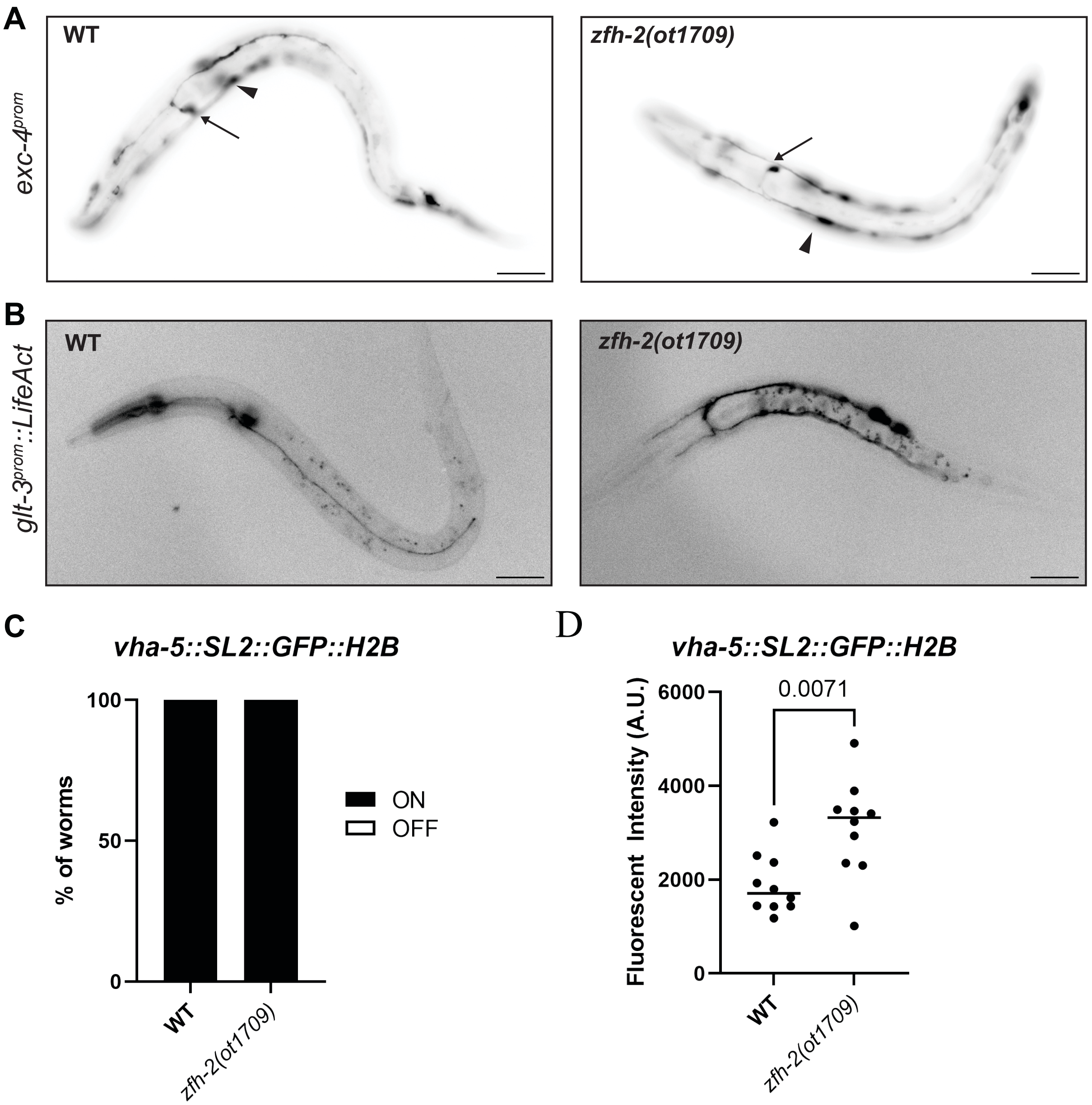


**Supplementary Figure S3: *zfh-2* null mutants show no obvious defects in the excretory cell (A)** An extrachromosomal array carrying an *exc-4prom::GFP* promoter fusion construct (*otEx669[exc-4p::GFP + rol-6(su1006)]*) in wild type and *zfh-2* null animals shows normal excretory canal morphology. Arrow is canal, arrowhead is hypodermal cell expression. **(B)** When expressed under a *glt-3* excretory cell specific promoter fragment (*arIs195[glt-3p::LifeAct::TagRFP]*), the actin reporter *LifeAct::TagRFP* shows similar excretory canal morphology in wild type and *zfh-2* null animals. **(C)** Animals homozygous for *zfh-2* null alleles retain expression of the excretory cell marker *vha-5(syb6835[vha-5::sl2::gfp::h2b])*. **(D)** Expression strength of the excretory cell marker *vha-5(syb6835[vha-5::sl2::gfp::h2b])* is not decreased in animals homozygous for *zfh-2* alleles.

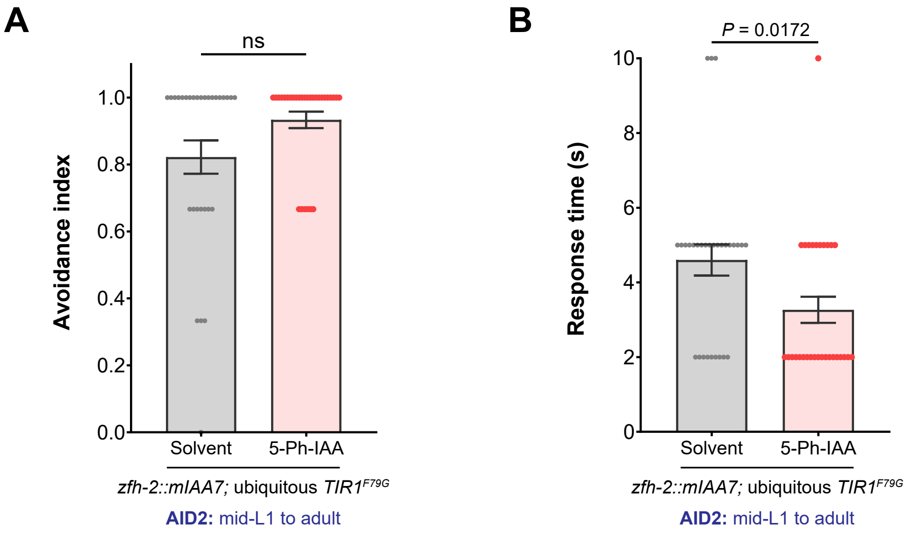


**Supplementary Figure S4: Postembryonic removal of ZFH-2 protein does not affect octanol avoidance behavior**
Avoidance index **(A)** and response time **(B)** of *zfh-2(syb10278); osIs158[eft-3p::TIR1(F79G)]* adults to 100% octanol. Animals were treated with either solvent (ethanol) or 100 µM 5-Ph-IAA starting at mid-L1 stage. Bars represent mean ± SEM. Data points represent individual animals. Denoted *P* value and not significant (ns) are from Mann-Whitney tests.

**Table S1: Expression pattern of ZFH-2::GFP protein in late larval/young adult stage animals.** Bold and underlined indicates that no homeobox identity regulator has yet been described for this neuron class. * indicates that expression in this neuron was previously overlooked.

| | | |
|---|---|---|
| Neurons | Anterior ganglion | IL1, OLQ, URY |
| | Other head ganglia | ADE, **ADF**, AFD, AIB, AIM, AIY, AIZ, ALA, ASG, ASH, **ASJ**, AVA, AVE, AVJ, AVL, AWB, CEP, FLP, RID, RIG, **RIM**, RIR, RIV, RMD, RMH, SAA, SAB, SIA, SIB, SMB, SMD |
| | Midbody | BDU, PDE, PVD, PVM, CAN* |
| | VNC | AS, DA, DB, VA, VB |
| | Tail | **DVA**, DVC, LUA, PDA, PDB, PHA, PHB, PQR, PVN |
| | Pharynx | NSM |
| Glia | All sheath glia | |
| | All socket glia | |
| | Not in GLR glia | |
| Excretory system | Canal cell | |
| | Gland cells | |
| | Duct cell | |
| | Pore cell | |
| Gonad | Spermatheca | |
| | Spermatheca-uterine valve | |
| | Gonadal sheath cells | |
| Vulva | Subset of vulval cells (vulE, vulF) | |
| Alimentary tract | Subset of pharyngeal muscle cells (pm1, pm2, pm7V, pm8) | |
| | Subset of pharyngeal epithelial cells (e1) | |
| | Subset of pharyngeal neurons (NSM only) | |
| | Pharyngeal-intestinal valve (VPI) | |
| | Not in hmc | |
| | Intestine: posterior intestinal cells only | |
| | Rectal valve cells (virL, virR) | |
| | Rectal gland cells (rect_D, rect_VL, rect_VR) | |
| | Rectal epithelial cells (U, Y, B, F, K, K') | |
| Body wall muscle | Not expressed | |
| Hypodermis | Not expressed | |

**Supplementary Table S2: Strain list.**

| **Strain** | ***Mutant or knock-in*** | **Array** | **DNA on array** | **Reference** |
|---|---|---|---|---|
| OH20426 | zfh-2(ot1812[zfh-2::gfp::loxP][*st12167]) I | | | This study |
| OH20145 | zfh-2(ot1709)/tmC20[unc-14(tmIs1219) dpy-5(tm9715)] I | | | This study |
| EG4887 | unc-119(ed3) III | oxIs322 | myo-2p::mCherry::H2B, myo-3p::mCherry::H2B, Cbr-unc-119(+) | PMID: 18953339 |
| OH15814 | him-5(e1490) V; dmd-4(ot935[dmd-4::gfp]) X | | | PMID: 33021200 |
| PHX4257 | eat-4(syb4257[eat-4::T2A::GFP::H2B]) III | | | PMID: 34415309 |
| PHX4491 | unc-17(syb4491[unc-17::T2A::GFP:H2B]) IV | | | PMID: 35324425 |
| PHX5463 | ins-6(syb5463[ins-6::SL2::gfp::H2B]) II | | | PMID: 36178933 |
| PHX6486 | cat-1(syb6486[cat-1::SL2::gfp::H2B]) X | | | PMID: 39422452 |
| PHX8255 | cat-2(syb8255[cat-2::sl2::gfp::h2b]) II | | | PMID: 39422452 |
| PHX6451 | tph-1(syb6451[tph-1::SL2::gfp::H2B]) II. | | | PMID: 39422452 |
| PHX7768 | tdc-1(syb7768[tdc-1::sl2::gfp::h2b]) II | | | PMID: 39422452 |
| OH15623 | | otIs706 | nlp-12p::tagRFPintrons | This study |
| PHX10278 | zfh-2(syb10278[zfh-2::mIAA7::wrmScarlet-I3::mIAA7]) I | | | This study |
| HS3528 | | osIs158 II | eft-3p::TIR1(F79G)::mRuby | PMID: 34865044 |
| OH19717 | | otIs935 V | UPNp:TIR1(F79G)::mTurquoise2:tbb-2 3' UTR, unc-122p:mCherry:unc-54 3' UTR | This study |
| OH20419 | zfh-2(syb10278[zfh-2::mIAA7::wrmScarlet-I3::mIAA7]) I | otSi4 II | myo-2p::TIR1(F79G)::mRuby::unc-54 3'UTR *ieSi60 | This study |
| FX17732 | zfh-2(tm310)/hT2[bli-4(e937) let-?(q782) qIs48] | | | National BioResource Project |
| FX22720 | zfh-2(tm12720) I | | | National BioResource Project |
| OH20425 | zfh-2(syb10278 ot1787[zfh-2$^{\Delta HD2}$])/tmC20[unc-14(tmIs1219) dpy-5(tm9715)] I | | | This study |
| OH20490 | zfh-2(syb10278 ot1797[zfh-2$^{\Delta HD3}$]) I | | | This study |
| OH20491 | zfh-2(syb10278 ot1787[zfh-2$^{\Delta HD2}$] ot1798[zfh-2$^{\Delta HD3}$])/tmC20[unc-14(tmIs1219) dpy-5(tm9715)] I | | | This study |
| OH20517 | zfh-2(syb10278 ot1807[zfh-2a M133>STOP] ot1808[zfh-2a M233>STOP M234>STOP) I | | | This study |

| | | | | |
|---|---|---|---|---|
| OH20518 | *zfh-2(syb10278 ot1809[zfh-2d M50>STOP]) / tmC20[unc-14(tmIs1219) dpy-5(tm9715)] I* | | | This study |
| OH17760 | | *otIs870* | *mir-228p::3XNLS::TagRFP* | PMID: 18085825, 39422452 |
| OH17400 | | *otIs857* | *mam-5prom2::3xNLS-TagRFP-T* | PMID: 32696701, 40950173 |
| PHX6670 | *spig-2(syb6670[spig-2::sl2::gfp::h2b]) V* | | | PMID: 38304163 |
| CM1360 | | *guIs30* | *clh-4::gfp* | PMID: 19921263 |
| QN305 | | *arIs195* | *glt-3p::LifeAct::TagRFP* | PMID: 27697907 |
| OH1277 | | *otEx669* | *exc-4::gfp; rol-6(su1006)* | PMID: 14684823 |
| AG408 | *pezo-1(av146[gfp::pezo-1]) IV* | | | PMID: 32490809 |
